# Phylogenomic Analyses Of 2,786 Genes In 158 Lineages Support a Root of The Eukaryotic Tree of Life Between Opisthokonts (Animals, Fungi and Their Microbial Relatives) and All Other Lineages

**DOI:** 10.1101/2021.02.26.433005

**Authors:** Mario A. Cerón-Romero, Miguel M. Fonseca, Leonardo de Oliveira Martins, David Posada, Laura A. Katz

## Abstract

Advances in phylogenetics and high-throughput sequencing have allowed the reconstruction of deep phylogenetic relationships in the evolution of eukaryotes. Yet, the root of the eukaryotic tree of life remains elusive. The most popular hypothesis in textbooks and reviews is a root between Unikonta (Opisthokonta + Amoebozoa) and Bikonta (all other eukaryotes), which emerged from analyses of a single gene fusion. Subsequent highly-cited studies based on concatenation of genes supported this hypothesis with some variations or proposed a root within Excavata. However, concatenation of genes neither considers phylogenetically-informative events (i.e. gene duplications and losses), nor provides an estimate of the root. A more recent study using gene tree-species tree reconciliation methods suggested the root lies between Opisthokonta and all other eukaryotes, but only including 59 taxa and 20 genes. Here we apply a gene tree – species tree reconciliation approach to a gene-rich and taxon-rich dataset (i.e. 2,786 gene families from two sets of ~158 diverse eukaryotic lineages) to assess the root, and we iterate each analysis 100 times to quantify tree space uncertainty. We estimate a root between Fungi and all other eukaryotes, or between Opisthokonta and all other eukaryotes, and reject alternative roots from the literature. Based on further analysis of genome size we propose Opisthokonta + others as the most likely root.

**Impact summary:** Finding the root of the eukaryotic tree of life is critical for the field of comparative biology as it allows us to understand the timing and mode of evolution of characters across the evolutionary history of eukaryotes. But this is one of the most challenging questions in evolutionary biology because the age (~1.8 billion years), diversity and complexity of eukaryotes challenge phylogenomic methods. This study evaluates the root using reconciliation of gene trees and species trees instead of the more common approach of analyzing concatenated genes. The dataset used in this study is bigger and more taxonomically inclusive than the dataset of any previous study about the eukaryotic root, and supports a root at or within Opisthokonta (e.g. animals and fungi). Finally, we explicitly test alternative hypotheses from the literature, and again find support for an Opisthokonta root.

## Introduction

One of the most controversial topics in the study of the history of life on Earth is the location of the root of the eukaryotic tree of life (EToL), which likely dates to around 1.6-1.8 billion years ago (de Duve 2007; Parfrey, et al. 2011). While there has been substantial progress on defining major eukaryotic clades such as Archaeplastida, Opisthokonta, SAR and Amoebozoa (Rodriguez-Ezpeleta, et al. 2005; Steenkamp, et al. 2006; Burki, et al. 2007; Hampl, et al. 2009; Adl, et al. 2012; Jackson and Reyes-Prieto 2014; Cavalier-Smith, et al. 2015; Katz and Grant 2015), the location of the root of EToL remains elusive.

Amongst the most highly-cited hypotheses, a root within Archezoa (Cavalier-Smith 1989, 1993) or between Unikonta - Bikonta (Stechmann and Cavalier-Smith 2002, 2003; Derelle and Lang 2011; Derelle, et al. 2015) have been proposed. The now-falsified Archezoa root proposed amitochondriate eukaryotes (e.g. microsporidians, diplomonads (e.g. *Giardia*), parabasalids (e.g. *Trichomonas*)) as the earliest-diverging lineages with all other mitochondria-containing lineages radiating after this divergence. This hypothesis lost support when the lack of mitochondria was demonstrated to be a derived character (Roger 1999).

In the past two decades, the Unikonta - Bikonta root has gained popularity and can be found in many text books. Though both clades have incorporated numerous taxonomic changes over the years, the root was first articulated as being between Opisthokonta + Amoebozoa and the rest of the eukaryotes based on a unique gene fusion event between dihydrofolate reductase (DHFR) and thymidylate synthase (TS) genes (Stechmann and Cavalier-Smith 2003). More recently, a new clade including Unikonta and former bikont lineages (i.e. Apusozoa, Breviata) was defined as Amorphea (Adl, et al. 2012) with the root dividing Amorphea and the remaining eukaryotes (Derelle, et al. 2015).

Advances in high-throughput sequencing technologies allow better estimation of eukaryotic phylogeny by providing the opportunity to explore bigger datasets and include non-model organisms such as rhizarians *Quinqueloculina* or the glaucophyte *Gloeochaete* (Burki, et al. 2007; Jackson and Reyes-Prieto 2014; Katz and Grant 2015; Brown, et al. 2018). A popular approach to take advantage of such opportunities is by inferring phylogenies from supermatrices by concatenating multiple genes in a single alignment (Rodriguez-Ezpeleta, et al. 2005; Dunn, et al. 2008; Wickett, et al. 2014; Derelle, et al. 2015). Analyses of multiple concatenated eukaryotic genes of putatively bacterial origin (i.e. mitochondrial) have either supported the Unikonta-Bikonta root (Derelle and Lang 2011; Derelle, et al. 2015) or suggested a new root between Discoba (Excavata) and the other eukaryotes (He, et al. 2014). Alternative methods have supported diverse root possibilities. For instance, a genome-wide analysis of rare genomic changes suggests a root between Archaeplastida and the other eukaryotes (Rogozin, et al. 2009), and an analysis based on the presence/absence of an encounter structure for the endoplasmic reticulum and the mitochondria suggests a root between Amorphea + Excavata and the rest of eukaryotes (Wideman, et al. 2013).

Phylogenomic methods vary in their approach to identify and account for evolutionary events such as lateral gene transfer (LGT), gene transfer from endosymbiosis and gene duplications/losses, which can be prevalent in many eukaryotic lineages (Galtier and Daubin 2008; Burki, et al. 2014; Katz 2015; Panchy, et al. 2016). Supermatrix methods require identifying paralog sequences and merging them into orthologous groups before building the concatenated alignment. Yet, distinguishing orthology from paralogy can be very difficult, particularly at scales of >1 billion years of eukaryotic evolution, and implicitly rely on knowledge of the duplication events. Despite the limitations of supermatrix methods, which discard informative data (e.g. gene duplications and losses), their tractability has made them popular choices in studies estimating the root of EToL.

There are also alternative methods that estimate the species tree from a set of gene trees. In contrast to supermatrix methods, these gene tree – species tree reconciliation methods incorporate informative data from different evolutionary events (e.g. gene duplication, LGT). These methods were designed to explore discordance between gene trees and species tree due to either incomplete lineage sorting (Mirarab, et al. 2014; Mirarab and Warnow 2015), gene duplication and loss (Chaudhary, et al. 2010) or LGT (Whidden, et al. 2014). One method for species tree inference is gene tree parsimony (GTP), which not only takes advantage of the power of gene-rich databases but can also consider disagreements across individual gene trees due to gene duplication, gene loss, LGT or/and incomplete lineage sorting. Other methods are based on more or less explicit models of gene tree evolution within species trees (De Oliveira Martins, et al. 2016; Mallo and Posada 2016), which in general substantially increase the needs for computational power.

Few studies have used reconciliation methods to estimate a rooted EToL. Based on only 20 gene trees, a preliminary GTP analysis estimated a root between Opisthokonta and the rest of eukaryotes (Katz, et al. 2012), which is consistent with initial analysis of a gene fusion (Stechmann and Cavalier-Smith 2002). Here we leverage a much larger dataset applying GTP to infer a rooted EToL. For this purpose, we used the recently published phylogenomic pipeline PhyloToL (Ceron-Romero, et al. 2019) and built a database of phylogenetic trees from 2,786 gene families including up to 158 species distributed across the whole EToL. To reduce the risk of the being trapped in local optima during heuristic search of gene tree rooting and of initial starting species tree generation and to better explore the tree space, we iterated analyses 100 times using four different datasets. We further evaluated and compared the levels of support for the most relevant previously published hypotheses.

## Methods

### Taxa selection

We started with the database of our phylogenomic pipeline PhyloToL (Grant and Katz 2014; Ceron-Romero, et al. 2019), which contains 1,007 taxa including Bacteria, Archaea and Eukaryotes. From this database, we generated two subsets of 158 and 155 eukaryotic taxa with two different criteria: 1) selecting taxa based on the quality of the data and maximizing the diversity based on their taxonomy (SEL+) and 2) selecting taxa randomly among the major eukaryotic clades Opisthokonta, Amoebozoa, Archaeplastida, Excavata, SAR and some orphan lineages (RAN+; Table 1 and Table S1). We also generated two extra datasets without microsporidians (SEL- and RAN-) in order to account on a possible effect over the phylogenetic inferences due to microsporidians fast-evolutionary rates.

**Table 1.**
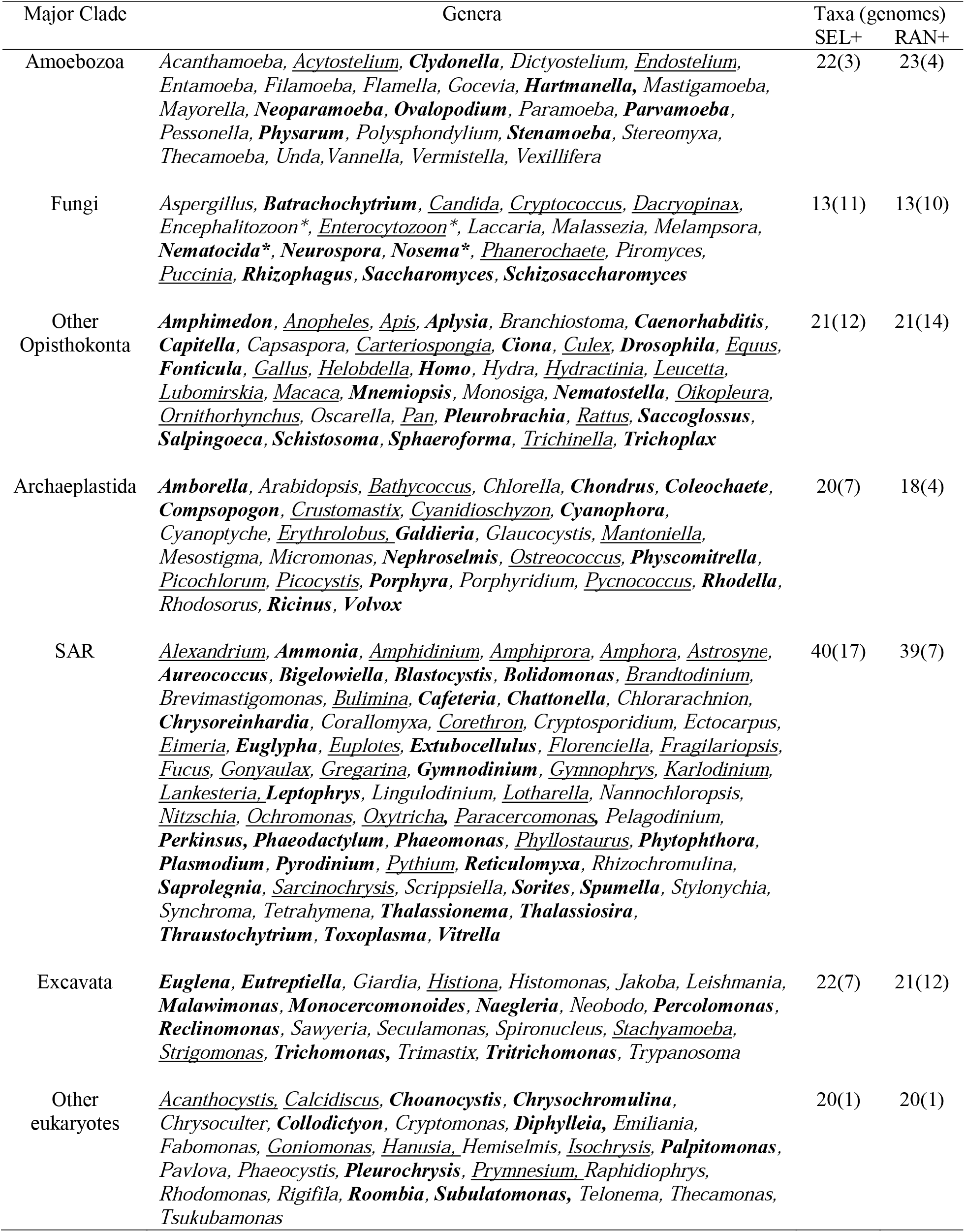
Summary of taxon selection for each study. Genera in **bold** are only in the taxonomy informed selected datasets (i.e., SEL+), <u>underlined</u> genera are only in the randomly selected within clades datasets (i.e, RAN+). The genera with asterisk (*) are microsporidians, which we excluded from datasets SEL- and RAN- because they often fall on very long branches (Embley and Hirt 1998; Hirt, et al. 1999; Van de Peer, et al. 2000). The numbers represent the amount of species included and the number of whole genomes in parenthesis (More details are in Table S1).

### Gene family selection

PhyloToL contains 13,104 protein-coding gene families. We chose the gene families that contain at least 25 taxa representing at least four of the five major eukaryotic clades. Additionally, at least 2 of the major clades had to contain at least 2 minor clades (e.g. Glaucophytes and Rhodophyta are minor clades in the major clade Archaeplastida). In a pilot analysis, we produced an alignment and a phylogenetic tree for each gene family using the default settings of a previous version of PhyloToL (GUIDANCE V1.3.1 sequence cutoff = 0.3 and column cutoff = 0.4; RAxML quick tree with model PROTGAMMALG and no bootstraps; Stamatakis 2006; Penn, et al. 2010; Grant and Katz 2014). Then, we kept the gene families that are exclusive of eukaryotes or the ones in which eukaryotes were monophyletic. From a total of 3,002 gene families that met our criteria, 2786 passed the initial steps of PhyloToL when including only the data from the dataset SEL+. This 2,786 gene families were used for further analyses with all datasets (Table S2).

### Multiple Sequence Alignments (MSA) and gene tree inference

MSAs for the four datasets were produced with PhyloToL (GUIDANCE V2.02 sequence cutoff = 0.3, column cutoff = 0.4, number of iterations = 5; Sela, et al. 2015; Ceron-Romero, et al. 2019). The default parameters of PhyloToL include up to five iterations of GUIDANCE V2.02 with 10 bootstraps and MAFFT V7 (Katoh and Standley 2013) with algorithm E-INS-i for less than 200 sequences or “auto” option if more than 200 sequences, and maxiterate = 1000. Instead, here we run the up to five iterations of GUIDANCE with 20 bootstraps and the simple MAFFT algorithm FFT-NS-2. Then, we perform an additional GUIDANCE run with 100 bootstraps and the default MAFFT parameters for PhyloToL.

Two approaches were applied for gene tree inference according to the method for inference of rooted species tree (i.e., *guenomu* and iGTP, see below). For *guenomu* (De Oliveira Martins, et al. 2016), gene trees were produced with MrBayes V3.2.1 (Huelsenbeck and Ronquist 2001; Ronquist, et al. 2012) using the dataset SEL+. The best-fit model of protein evolution was selected using ProtTest v.3.3 (Darriba, et al. 2011) based on the Bayesian Information Criterion. MrBayes was run with four Markov chains incrementally heated and started with a randomly generated tree. For every 100 generations of 1×10^7^ generations, we sampled one tree. The posterior distribution of trees, after discarding 25% as burn-in, was summarized by the trees comprising the 50% highest posterior set, to decrease the effect of trees sampled only once. For iGTP, gene trees were inferred with RAxML v.8.2.4 (Stamatakis 2014) with 10 ML searches for best-ML tree (option “-# 10”), using rapid hill-climbing algorithm (option “-f d”) and no bootstrap replicates. The protein evolution model used was evaluated during the gene tree inference (option “-m PROTCATAUTO”) by testing all models available in RAxML (e.g. JTT, LG, WAG, etc) with optimization of substitution rates and of site-specific evolutionary rates which were categorized into four distinct rate categories for greater computational efficiency.

### Inference of rooted species trees

In order to infer a rooted EToL, we set out to use two supertree tools for species tree inference, the Bayesian-based *guenomu* and the gene tree parsimony tool iGTP. While iGTP considers that the disagreement between gene trees and the species tree is due to either duplications, duplications-losses or deep coalescence; *guenomu* considers the effect of these and other evolutionary processes in a multivariate manner. With *guenomu* we did not see convergence in two independent replicates in a reasonable time, which may reflect a lack of convergence of MrBayes or underlying uncertainty in the gene trees; therefore, we chose to continue further analyses with iGTP only, which relies on point estimates of the gene phylogenies.

We ran iGTP for the four datasets with the analysis option that account for gene duplications and losses. In our application of iGTP, we decided to iterate each analysis 100 times to explore the tree space. Given the complexity of the datasets and the heuristic nature of some key steps of the iGTP algorithm (e.g. gene tree rooting and of initial starting species tree generation), in a preliminary analyses we faced two systematic challenges with iGTP as the inferred species tree was affected by: 1) the order of the leaves in the input unrooted gene tree newick strings (i.e. the input trees were treated as rooted even though we specified that they were not); and 2) the input gene order in the 100 replicates. Therefore, we randomly shuffled the order of the leaves in the unrooted gene trees (keeping the same topology), and randomly shuffled the order of the input gene trees in each of the 100 replicates per dataset.

### Comparing different EToL root hypotheses

For the datasets SEL+, RAN+, SEL- and RAN-, we compare 5 different hypotheses of the root of EToL. These hypotheses are: 1) the most parsimonious root according to the iGTP analysis, 2) between Opisthokonta and the rest of eukaryotes, 3) between Discoba (Excavata) and rest of the eukayotes, 4) between Unikonta and Bikonta, and 5) between Metamonada (Excavata) + Ancyromonadida and the rest of eukaryotes. For the Unikonta-Bikonta root, different alternative topologies were evaluated according to the multiple changes on the definition of the Unikonta clades, but only the one with the lowest reconciliation cost was used for further comparisons. In order to compare the hypotheses, we constrained species trees (fixing the relationships among major clades while allowing the relationships within major clades to be inferred by iGTP) according to every hypothesis and calculated the reconciliation cost per hypotheses in each dataset.

### Computational resources

The production of alignments following the strategy described above for each of the four datasets required 10 weeks of running time (around 40 weeks in total) in 75 threads and around 120 GB of RAM. The gene tree inference for each dataset required around 4 weeks (around 16 weeks in total) in 24 threads and 24 GB of RAM. Each iGTP analysis (with 100 replicates) requires 1 week of running time in 100 threads and ~100 GB of RAM. Given that there were 6 iGTP analyses per dataset, the running time for all datasets was around 24 weeks.

### Quantification and statistical analysis

We iterated every iGTP analysis 100 times to quantify tree space uncertainty. Given this large sample size (n=100), we decided to use Kolmogorov–Smirnov and a significance level of 0.01 to test if the reconciliation costs of each root hypothesis distributed as normal. Then, we compared the mean reconciliation costs between Fungi + others and every other hypothesis in all datasets using t-tests. The results of these tests and measures of mean and standard deviation were summarized in Tables S3 and S4 and also displayed in Figure 2.

## Results

### Building the phylogenomic datasets

We used the database of PhyloToL, which contains 13,104 gene families and 1,007 taxa (i.e. including Archaea, Bacteria and Eukaryotes; Grant and Katz 2014; Ceron-Romero, et al. 2019) to select the gene families for this study. Initially, we filtered the gene families that were present in at least 25 taxa of at least 4 major eukaryotic clades (i.e. Opisthokonta, Amoebozoa, Archaeplastida, Excavata and SAR; Table 1; Table S1). Then, we built alignments and phylogenetic trees to select the 2,786 gene families that are only found in eukaryotes or in which eukaryotes are monophyletic (see 2. Material and methods; Table S2).

In order to balance phylogenetic diversity and computation speed, we built four datasets that each included the 2,786 aforementioned gene families and up to 158 eukaryotic species from between 140 and 158 genera (Table 1; Table S1 and S2). The four datasets varied based on taxon selection criteria: for the ‘SEL+’ (i.e. selected) dataset, we selected representative species of all major eukaryotic clades based on our assessment of data quality and taxonomic breadth; and for the ‘RAN+’ (i.e. random) dataset, we randomly chose even numbers of species among the major eukaryotic clades. We also generated two additional databases by excluding the fast-evolving Microsporidia (i.e. SEL- and RAN-) as the inclusion of these lineages can generate phylogenetic artifacts such as long-branch attraction (Embley and Hirt 1998; Hirt, et al. 1999; Van de Peer, et al. 2000).

### Inference on location of EToL root

Though we set out to deploy two gene tree – species tree reconciliation methods to infer the root of the eukaryotic tree of life, due to the complexity of the data, we were constrained to focus on only one method for the analyses presented here. Our original intent was to use both a Bayesian supertree approach with the software *guenomu* (De Oliveira Martins, et al. 2016) and a gene tree parsimony approach with the software package iGTP (Chaudhary, et al. 2010). Unfortunately, *guenomu* failed to converge in an estimate of species trees after being run for multiple weeks on a cluster with more than 400 cores, likely due to the complexity of the algorithm and underlying uncertainty in the gene trees, so we continued only with iGTP.

Using iGTP, we estimated the most parsimonious rooted tree of eukaryotes for each of our four datasets, all of which indicate Fungi as the earliest branching group (Figure 1). Other less parsimonious but frequent alternatives indicate glaucophytes or the apusozoan *Fabomonas tropica* as the earliest branching group or taxon. Across all replicates of the analysis, the second most frequent earliest branching group is Opisthokonta (i.e. the remaining opisthokonts when the earliest branching group was Fungi). These results leave open the possibility of a root between Opisthokonta and the other eukaryotes, which we discuss below.

**Figure 1.**
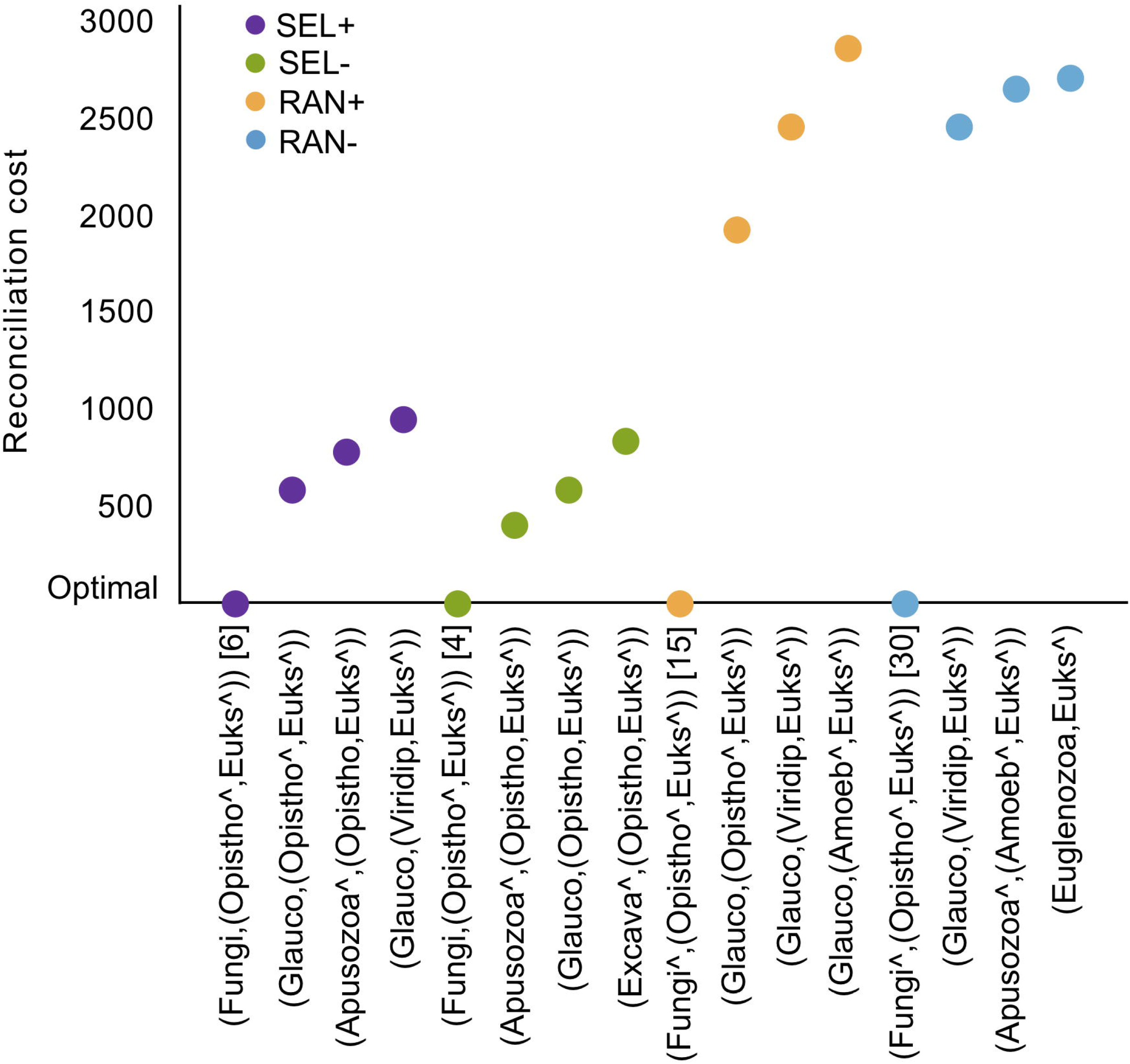
A root between fungi and all other eukaryotes is the most parsimonious hypothesis based on 100 iterations of iGTP using all four taxon sets. Here we report the four most parsimonious topologies in 100 iterations of the analysis, and note the number of times the first hypothesis appeared before any alternative in square parenthesis (i.e. a fungal root was present in the six iterations of iGTP with lowest reconciliation scores in the SEL+ analyses). SEL+: Taxonomically informed taxa selection including microsporidians, SEL-: taxonomically informed taxa selection excluding the long-branch microsporidians, RAN+: random taxa selection including microsporidians, RAN-: random taxa selection excluding microsporidians. The caret (^) implies a non-monophyletic clade. For example, in datasets SEL+ and RAN+ the microsporidians do not fall in the same clade as the rest of opisthokonts. We show the relative reconciliation costs compared to optimum (lowest value) for each dataset. After fungi + other eukaryotes, other parsimonious roots involve underrepresented clades in our database such as Glaucophytes or some apusozoans (see also Figure S2).

### Comparison to published EToL hypotheses

We also used iGTP to evaluate various hypotheses from the literature including a root between Opisthokonta and others (Stechmann and Cavalier-Smith 2002; Katz, et al. 2012), between Discoba (Excavata) and others (He, et al. 2014), the Unikonta – Bikonta root (Stechmann and Cavalier-Smith 2003; Derelle, et al. 2015). Additionally, we included an alternative root with Ancyromonadida + Metamonada as sister to all others eukaryotes (personal communication Tom Williams, Celine Petitjean), which emerged from studies of probabilistic gene tree-species tree reconciliation analyses with amalgamated likelihood estimation (ALE; Szollosi, Rosikiewicz, et al. 2013; Szollosi, Tannier, et al. 2013). Here, iGTP estimates the reconciliation cost of a species tree given constrained phylogenetic relations among major eukaryotic clades to reflect the different hypotheses of the root of EToL (Figure 2, x-axis). In addition to these four hypotheses, we also calculated and compared the reconciliation cost of a species tree reflecting our initial estimate, placing the root between Fungi and the other eukaryotes. The results show that for the datasets SEL- and RAN- our inferred root of Fungi + others is more parsimonious than the other 4 hypotheses, while for dataset SEL+ and RAN+ the most parsimonious root is Opisthokonta + others (Figure 2).

**Figure 2.**
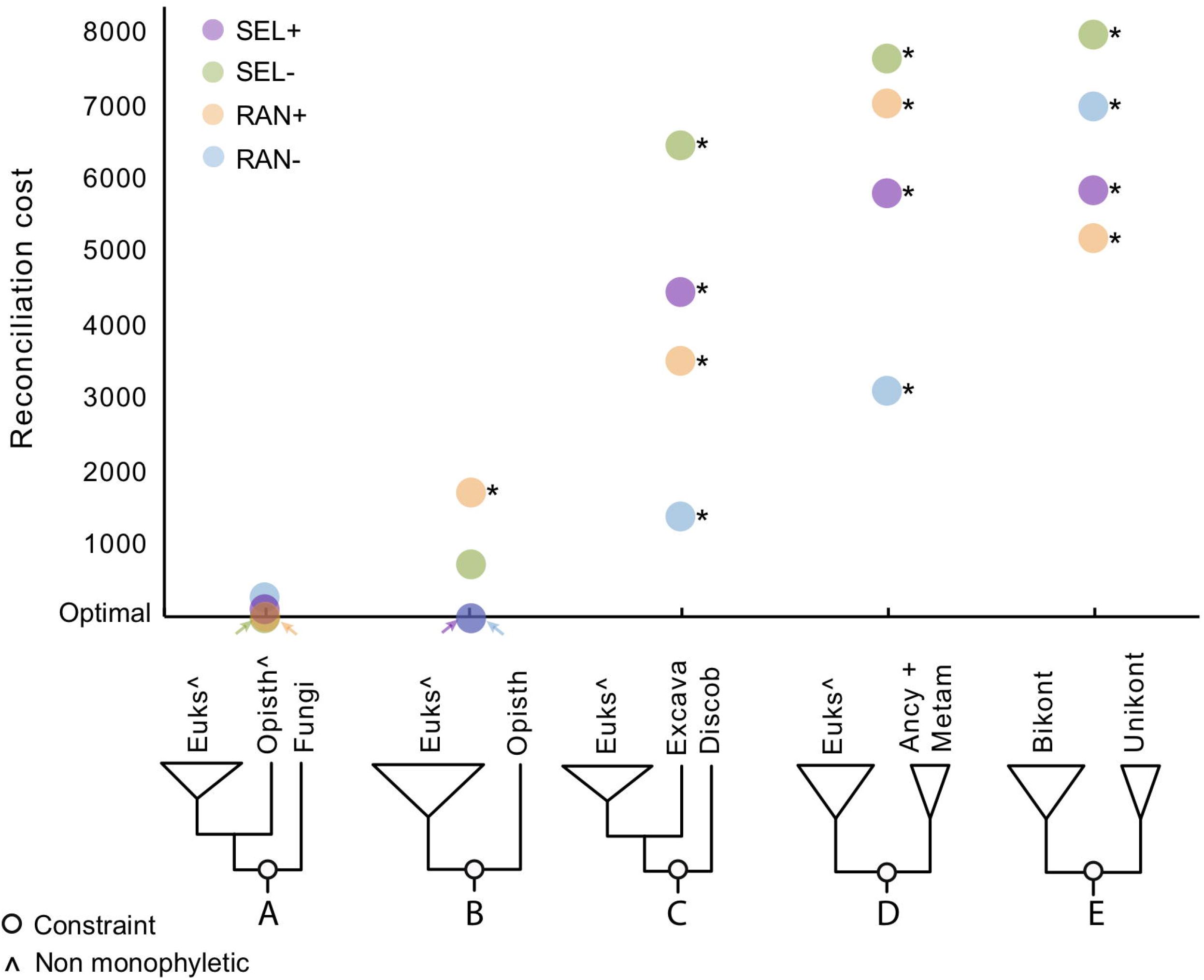
Constraining the species tree to match varying hypotheses of the root of EToL supports a root at or within Opisthokonta and is inconsistent with other hypotheses. We show the relative reconciliation costs compared to optimum (lowest value) for each dataset. The five hypotheses here are: A) between fungi and other eukaryotes (our estimate from the previous analysis), B) between Opisthokonta and other eukaryotes (Stechmann and Cavalier-Smith 2002; Katz, et al. 2012), C) between Ancyromonadida + Metamonada and the others, D) between Discoba and the others (He, et al. 2014), and E) between unikonta and bikonta (Stechmann and Cavalier-Smith 2002; Derelle, et al. 2015). The empty circles on the cartoon phylogenies indicate where in the tree the constraint was applied, and other notations are as in Figure 1. Overall, the reconciliation costs for Fungi + others is similar to the reconciliation cost of Opisthokonta + others and significantly different (asterisks) to the reconciliation costs in all other hypotheses (t-student, p > 0.001; Tables S3 and S4), except in the database RAN+. This result may imply that Opithokonta + other is the root while Fungi + others could be a potential artifact.

We assessed the difference in reconciliation costs between Fungi + others and every other hypothesis in all datasets. We determined that reconciliation cost values are normally distributed based on QQ plots and Kolmogorov–Smirnov tests (n = 100, p > 0.01; Table S3). Then, we performed t-tests to determine if there is a statistically significant difference between the mean reconciliation costs of the Fungi + others hypothesis against every other hypothesis. Our results show that for datasets SEL+, SEL- and RAN+ there are no significant statistical differences between the mean reconciliation costs of Opisthokonta + others and Fungi + others (n = 100, p > 0.01; Tables S3 and S4); for the RAN-, the mean reconciliation cost for the rooted species tree (Fungi + others) is significantly lower than (Opisthokonta + others). For all four taxon sets, the mean estimated reconciliation costs for species trees inferred to match the remaining published hypotheses are all statistically significantly higher (i.e. less parsimonious) than our rooted species tree (Fungi + others, t-student, p < 0.01; Figure 2 and Tables S3 and S4).

### Assessing the effect of missing data in Fungi over root calculations

We tested if missing data in fungi, due to reduced genomes (Figure S1), artifactually contribute to the most parsimonious root between Fungi/Opisthokonta and the rest of eukaryotes. We ran iGTP in two subsets from the SEL+ dataset: 1) 336 genes that contain at least 10 metazoan and 10 fungi species and 2) 246 genes that contain at least 10 metazoans and no fungi. For the first subset the most parsimonious root was between the apusozoan *Fabomonas tropica* (one of the taxa with more missing data; Fig S2 and Table S1) and the others, while the root between Fungi and the others was the second most parsimonious (Figure S3). In both topologies, the next earliest divergent group was *other* Opisthokonta. For the small set of genes present only in 10 or more metazoan and no fungi, a root between Opisthokonta and others still appears as one of the most parsimonious roots (Figure S4).

## Discussion

This study analyzed 2,786 gene trees for four taxon sets of up to 158 diverse eukaryotic taxa, each iterated 100 times by changing both gene tree order and root, which is perhaps the largest analysis yet to address the root of the eukaryotic tree of life. As in Katz, et al. (2012), we used gene tree parsimony as implemented in the software iGTP to estimate the root of EToL that minimizes gene duplications and loses. Given the importance of gene duplication/loss for the evolution of eukaryotic genomes (Wolfe and Shields 1997; Otto and Whitton 2000; Dehal and Boore 2005), their inclusion in the estimation of the most likely root of EToL represents a powerful alternative to studies that are based on a supermatrix approach, as the latter require users to discard paralogs that can contain phylogenetic information (Guigo, et al. 1996; Chaudhary, et al. 2010).

Across our analyses we find support for either Fungi or Opisthokonta as sister to all other eukaryotes (Figures 1 and 2). The Fungi + others root is consistently the most parsimonious across all taxon datasets (Figure 1). Martin, et al. (2003) argued for this hypothesis based on the fact that fungi have osmotrophic feeding while most other eukaryotes are phagotrophic (with the exceptions including autotrophic lineages). Moreover, fungi contain more ATP pathways than other eukaryotic clades, including the ability to synthesize ATP under anoxic and high sulfide conditions that may resemble the environment during early eukaryotic evolution (Martin, et al. 2003). Analyses of the fossil record provide evidence consistent with a hypothesis of ‘fungi first’ as some pre-Ediacaran fossils look similar to fungi, though more work needs to be done to robustly classify them (Butterfield 2005, 2009). Recent fossil finds in Arctic Canada are also consistent with a pre-Ediacaran origin of fungi (Loron, et al. 2019), and these fossils are twice as old as the fossil used for the current estimates of the origin of fungi (450 Ma; Redecker, et al. 2000). Although there are no previous phylogenetic studies supporting Fungi as the earliest branching eukaryotic clade and the monophyly of Opisthokonta is widely accepted (Baldauf and Palmer 1993), the fossil record findings together with the characteristics of energy production in fungi support further exploration of this hypothesis.

We explored an alternative root between Opisthokonta and all other eukaryotes as gene loss due to reductions in genome size (Braun, et al. 2000; Nagy, et al. 2014; Stajich 2017) and/or lateral gene transfer (LGT) from bacteria to fungi (Rosewich and Kistler 2000; Richards and Talbot 2013) may artificially pull fungi to the root. Estimating the reconciliation costs for trees constrained to have a root with either fungi or all of Opisthokonta as sister to other eukaryotes yielded similar results (Figure 2). A root between Opisthokonta and the other eukaryotes is consistent with initial analysis of a gene fusion (Stechmann and Cavalier-Smith 2002), though increasing taxon sampling indicated that these gene fusions had a more complex distribution (Stechmann and Cavalier-Smith 2003). A more recent study based on gene tree parsimony also proposed an Opisthokonta + others root (Katz, et al. 2012), but with a substantially smaller dataset than this study. Fungi have experienced high rates of gene loss (Braun, et al. 2000; Nagy, et al. 2014; Stajich 2017), which is reflected in their significantly reduced genome sizes when compared to animals (Figure S1). This outcome would be even more likely if Opisthokonta experienced frequent genome duplication events and many of the genes kept in Fungi came from different paralogs than the ones kept in the other Opisthokonta (Albalat and Canestro 2016; Fernandez and Gabaldon 2020; Guijarro-Clarke, et al. 2020). Similarly, several studies indicate that interdomain LGTs are frequent in fungi (Rosewich and Kistler 2000; Wenzl, et al. 2005; Lawrence, et al. 2011), and such genes might also pull fungi to the root of the eukaryotic tree of life.

Missing data may explain the unexpected support for a root of Glaucophyta + others (Glaucophyta, (Opisthokonta, others)), which appears as one of the four most parsimonious roots (though always with a higher reconciliation cost than Fungi + others) across taxon sets (Figure 1). Despite our effort on choosing genes with well-represented species, the glaucophytes are the minor clade with the fewest data across gene trees (Figure S2), likely due to incomplete sequencing of transcriptomes instead of high rates of gene loss. Previous studies have shown that the gene tree parsimony approach for species tree inference is sensitive to missing data (Burleigh, et al. 2011; Davis, et al. 2019). Because iGTP doesn’t distinguish gene loss from missing data, genome reduction from the completely sequenced fungal genomes and missing data in the poorly-sampled Glaucophyte can both lead to clades being pulled to the root.

It could also be argued that the large amount of ‘missing data’ within fungi, which is driven by relatively small genome sizes, creates an artifactual placement of the fungal/Opithokonta root. We tested for this effect in two analyses: 1) 336 genes present in ≥10 species of both fungi and metazoan, and 2) 246 genes present in >10 metazoa but absent from fungi. In both analyses, a root between Opisthokonta or Fungi and the others was still one of the most parsimonious roots (Figures S3 and S4). In contrast to analyses of all 2,786 genes, numerous hypotheses have very similar reconciliation costs, likely due to the reduced power of such small gene sets. Nevertheless, the retention of fungi/opisthokont root in both analyses suggests that ‘missing data’ in the fungi are not a determining factor for the results of the fuller analyses.

Analyses of the EToL that rely on a supermatrix approach (e.g. Derelle and Lang 2011; He, et al. 2014; Derelle, et al. 2015; Brown, et al. 2018) are likely to be particularly sensitive to challenges in ortholog selection with incomplete data and/or when inclusion of orphan lineages. Most of these studies are consistent with a Unikonta-Bikonta root, though with changing taxonomic definitions of the unikonts (e.g. Derelle and Lang 2011; Derelle, et al. 2015; Brown, et al. 2018). Reanalysis of a supermatrix study that originally proposed a root within Excavata (He, et al. 2014), similarly indicated a root between Opimoda (unikonts plus malawimonads and collodictyonids) and the rest of the eukaryotes (i.e. a revision of unikonta/biokonta root; Derelle, et al. 2015). The inclusion of preliminary transcriptome data from ancyromonads was also found to be consistent with the Unikonta-Bikonta root, placing ancyromonads in the Unikonta side and close to the root (Brown, et al. 2018). In these supermatrix analyses, selection of orthologs based on single tree topologies from incomplete datasets (i.e. incomplete transcriptome data) are likely to induce bias in the inference of the placement of the root. Intriguingly, studies that use methods other than a supermatrix approach predict roots other than the Unikonta-Bikonta (e.g. Stechmann and Cavalier-Smith 2002; Rogozin, et al. 2009; Katz, et al. 2012; Wideman, et al. 2013), again suggesting that the Unikonta-Bikonta root could be a biproduct of supermatrix analyses, which ignore phylogenetically-informative gene duplication events.

In conclusion, our estimates of the root of the eukaryotic tree of life based on 2,786 genes are consistent across taxon datasets and are robust when compared to other hypotheses in the literature. There are clearly caveats to be considered. For example, though we sought to remove genes that included interdomain LGTs (see 2. Material and methods), it is possible that we missed cases in our pilot analyses and as a result, examples of interdomain LGT ‘pull’ fungi to the root. Further, we acknowledge that missing data likely impacts our estimates, as we argue that the spurious position of a root between glaucophytes and other eukaryotes is due to lack of genome data. These two factors and the difference of genome size between metazoans and fungi support the idea that the root falls between Opisthokonta and the rest of the eukaryotes. Future studies with more complete genome data, ideally implemented with algorithms that explicitly account for gene duplications/losses and LGT, are required to validate the estimates presented here.

## Acknowledgements

We thank members of the Katz lab for comments on earlier versions of the manuscript. This work was supported by a National Institute of Health award (R15HG010409) and three awards from the National Science Foundation (OCE-1924570, DEB-1651908 and DEB-1541511) to L.A.K.

## Author contributions

LA.K and M.A.C.R conceived of the study and broad approach, and designed the experiments in collaboration with M.M.F and D.P. M.A.C.R, M.M.F and L.O.M performed the analyses. M.A.C.R and L.A. K wrote the manuscript with input from all authors.

## Data archiving

Raw and analyzed data are deposited in the DRYAD database (https://datadryad.org/stash/share/9jrGM0UhndWT0tYmoawBnwfkpB-cuNfnQwL9fPuxBiU)

## Declaration of Interests

None

## Notes

https://datadryad.org/stash/share/9jrGM0UhndWT0tYmoawBnwfkpB-cuNfnQwL9fPuxBiU

### Competing Interest Statement

The authors have declared no competing interest.

