## Supplementary material for "Phylogenomic Analyses Of 2,786 Genes In 158 Lineages Support a Root of The Eukaryotic Tree of Life Between Opisthokonts (Animals, Fungi and Their Microbial Relatives) and All Other Lineages": Table S3

**Table S3.** Normality test for the reconciliation costs of each root hypothesis in every dataset. Kolmogorov–Smirnov was chosen given the sample size n=100. The table shows the D statistic and the p-value in addition to the mean and standard deviation. Under a significance level of 0.01 all sets of replicates distribute as normal.

| dataset | H | mean | S. Dev | D | p-value |
|---|---|---|---|---|---|
| SEL- | Fungi | 578960 | 1853 | 0.126 | 0.082 |
| SEL- | Opisthokonta | 579504 | 1721 | 0.132 | 0.060 |
| SEL- | Unikonta | 586245 | 1000 | 0.067 | 0.757 |
| SEL- | Discoba | 584517 | 1011 | 0.107 | 0.204 |
| SEL- | Ancyromonadida + Metamonada | 586215 | 952 | 0.087 | 0.430 |
| SEL+ | Fungi | 585077 | 1748 | 0.161 | 0.011 |
| SEL+ | Opisthokonta | 584468 | 1605 | 0.124 | 0.093 |
| SEL+ | Unikonta | 589759 | 795 | 0.123 | 0.095 |
| SEL+ | Discoba | 588129 | 1227 | 0.136 | 0.050 |
| SEL+ | Ancyromonadida + Metamonada | 590196 | 1322 | 0.085 | 0.462 |
| RAN+ | Fungi | 393224 | 879 | 0.094 | 0.340 |
| RAN+ | Opisthokonta | 394779 | 1007 | 0.156 | 0.015 |
| RAN+ | Unikonta | 399810 | 610 | 0.069 | 0.725 |
| RAN+ | Discoba | 396328 | 689 | 0.102 | 0.253 |
| RAN+ | Ancyromonadida + Metamonada | 398050 | 594 | 0.108 | 0.194 |
| RAN- | Fungi | 398301 | 926 | 0.138 | 0.045 |
| RAN- | Opisthokonta | 398455 | 1060 | 0.108 | 0.198 |
| RAN- | Unikonta | 402879 | 579 | 0.083 | 0.492 |
| RAN- | Discoba | 399246 | 707 | 0.105 | 0.222 |
| RAN- | Ancyromonadida + Metamonada | 401042 | 762 | 0.113 | 0.154 |
