## Supplementary material for "Phylogenomic Analyses Of 2,786 Genes In 158 Lineages Support a Root of The Eukaryotic Tree of Life Between Opisthokonts (Animals, Fungi and Their Microbial Relatives) and All Other Lineages": Table S4

**Table S4.** Statistical comparison of the mean reconciliation costs between Fungi + others and every other hypothesis in all datasets. For each comparison, the table contains the T statistic, degrees of freedom (df) and the p-value.

| dataset | H1 | H2 | T | df | p-value |
|---|---|---|---|---|---|
| SEL- | Fungi | Opisthokonta | -2.154 | 196.940 | 0.032 |
| SEL- | Fungi | Unikonta | -34.607 | 152.140 | < 0.01 |
| SEL- | Fungi | Discoba | -26.327 | 153.200 | < 0.01 |
| SEL- | Fungi | Ancyromonadida + Metamonada | -34.837 | 147.840 | < 0.01 |
| SEL+ | Fungi | Opisthokonta | 2.564 | 196.570 | 0.011 |
| SEL+ | Fungi | Unikonta | -24.378 | 138.260 | < 0.01 |
| SEL+ | Fungi | Discoba | -14.292 | 177.480 | < 0.01 |
| SEL+ | Fungi | Ancyromonadida + Metamonada | -23.354 | 184.330 | < 0.01 |
| RAN+ | Fungi | Opisthokonta | -1.096 | 194.480 | 0.274 |
| RAN+ | Fungi | Unikonta | -41.909 | 166.180 | < 0.01 |
| RAN+ | Fungi | Discoba | -8.113 | 185.130 | < 0.01 |
| RAN+ | Fungi | Ancyromonadida + Metamonada | -22.863 | 190.890 | < 0.01 |
| RAN- | Fungi | Opisthokonta | -11.636 | 194.450 | < 0.01 |
| RAN- | Fungi | Unikonta | -61.562 | 176.410 | < 0.01 |
| RAN- | Fungi | Discoba | -27.788 | 187.370 | < 0.01 |
| RAN- | Fungi | Ancyromonadida + Metamonada | -45.486 | 173.900 | < 0.01 |
