## Supplementary material for "Phylogenomic Analyses Of 2,786 Genes In 158 Lineages Support a Root of The Eukaryotic Tree of Life Between Opisthokonts (Animals, Fungi and Their Microbial Relatives) and All Other Lineages": Figure S1

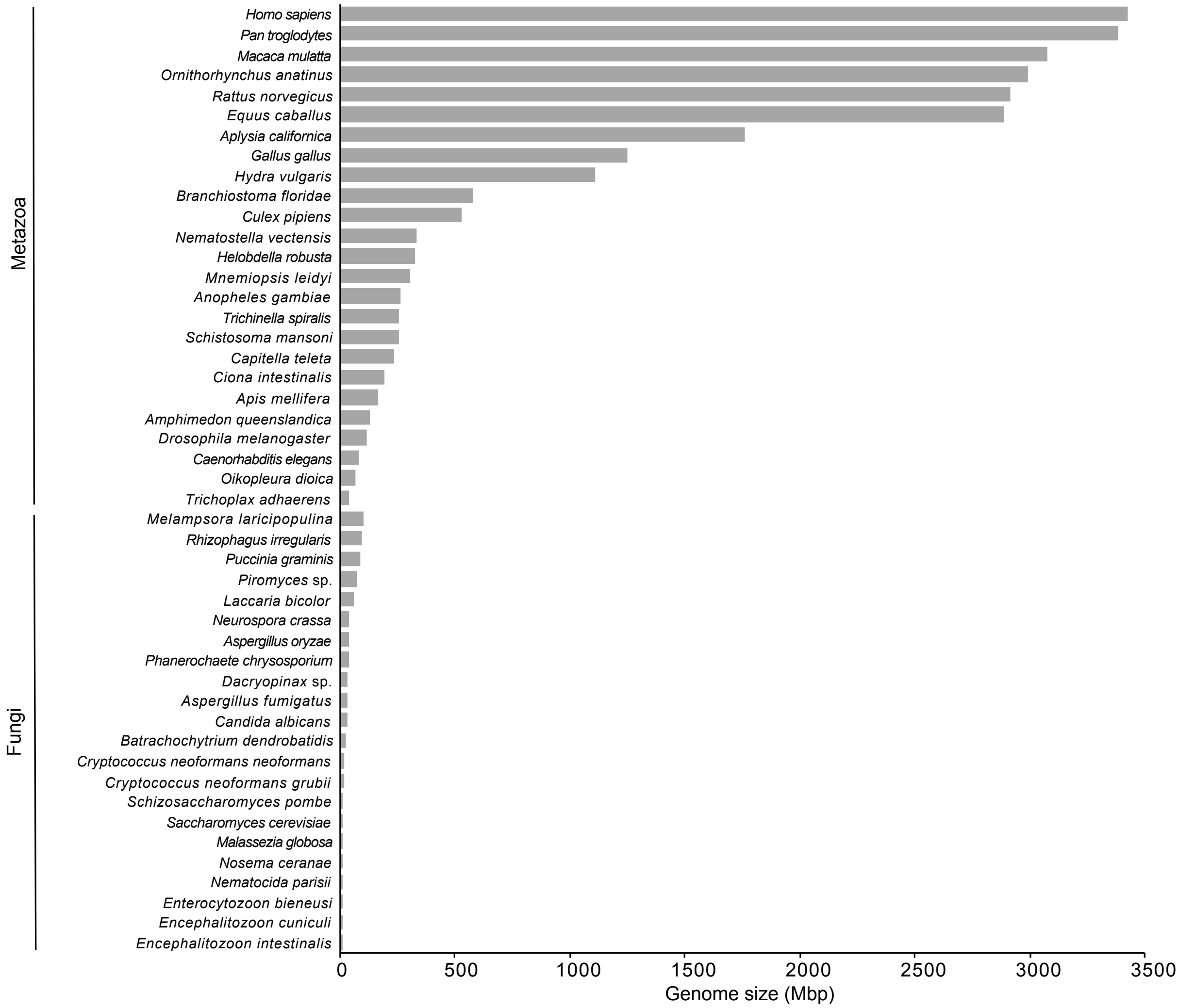


**Figure S1.** Genome size comparison between Metazoa and Fungi. The type of data used for these taxa were 25 whole genomes and 7 transcriptomes for metazoans and 21 genomes and 1 transcriptome for fungi (Table S1). The fungi genome sizes were taken from JGI (https://jgi.doe.gov/) and the metazoan genome sizes were taken from the Animal Genome Size Database, Release 2.0 (http://www.genomesize.com).
