## Supplementary material for "Phylogenomic Analyses Of 2,786 Genes In 158 Lineages Support a Root of The Eukaryotic Tree of Life Between Opisthokonts (Animals, Fungi and Their Microbial Relatives) and All Other Lineages": Figure S2

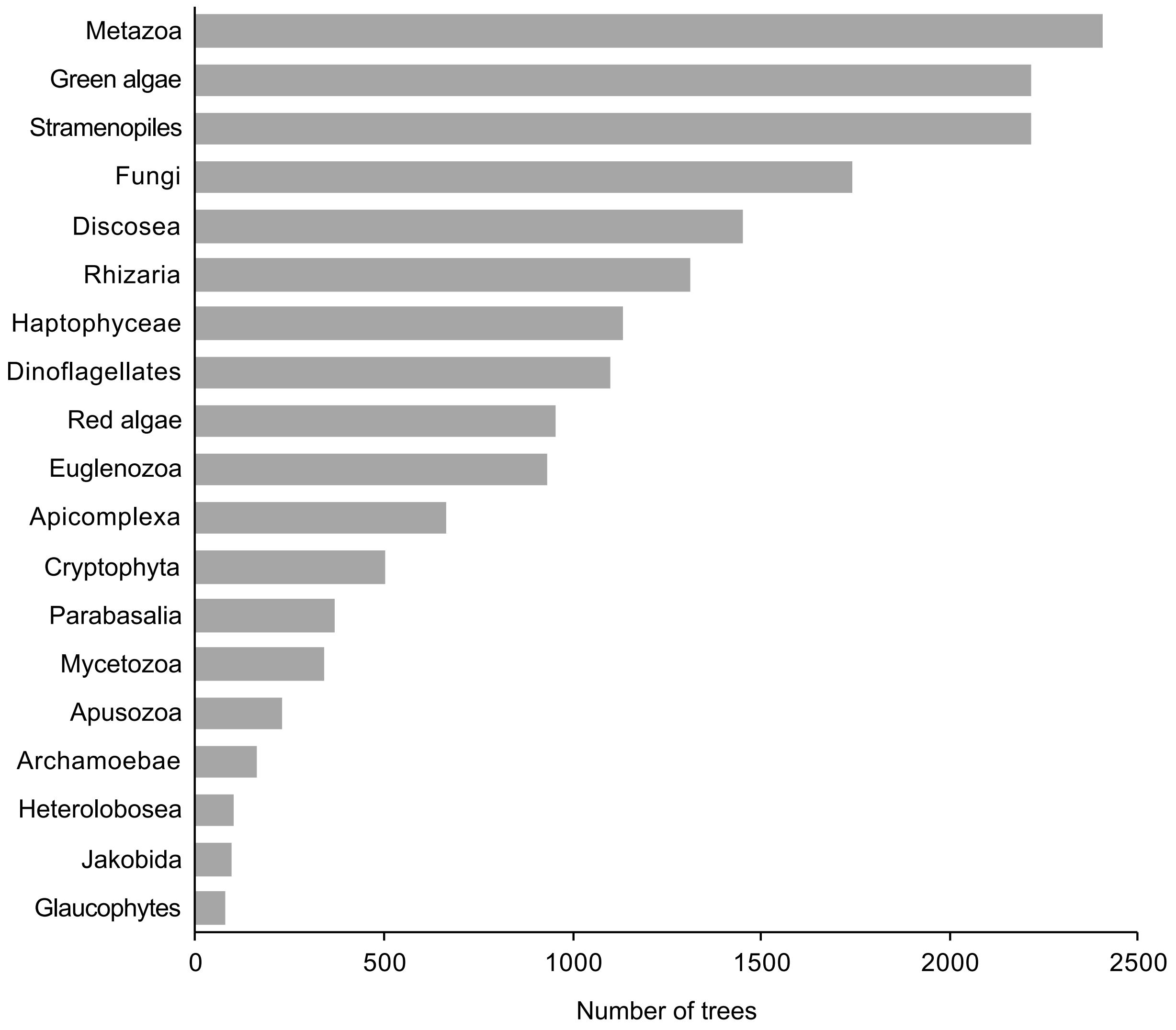


**Figure S2.** Number of trees with at least three species per minor clade in dataset SEL+. The data used for all these clades were a combination of whole genome sequences and transcriptomes. For Glaucophytes, the most underrepresented clade in the trees, all data (three taxa; Table S1) came from transcriptomes. The lack of data in Glaucophytes, some excavate clades and Apusozoa may affect in some cases the root assessments by iGTP.
