## Supplementary material for "Phylogenomic Analyses Of 2,786 Genes In 158 Lineages Support a Root of The Eukaryotic Tree of Life Between Opisthokonts (Animals, Fungi and Their Microbial Relatives) and All Other Lineages": Figure S3

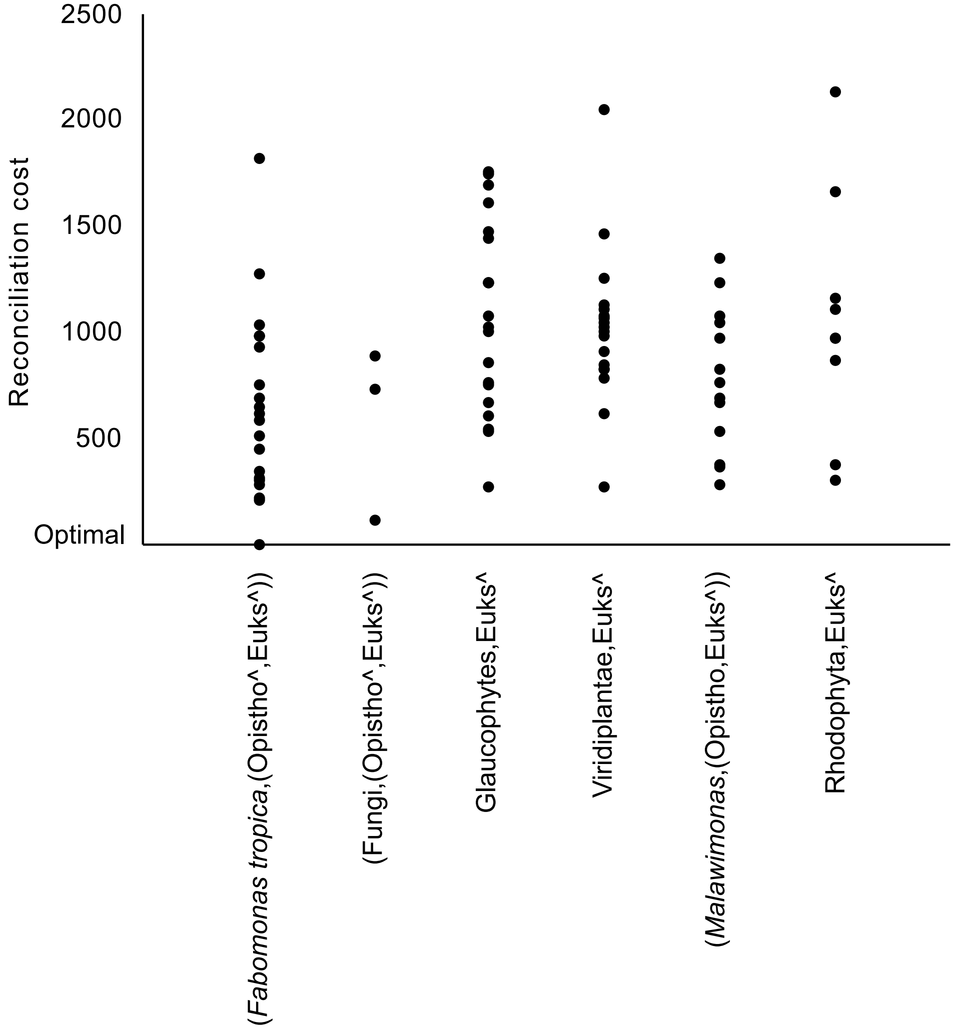


**Figure S3.** To assess whether the reduced gene numbers within fungal genomes drives our estimate of the root, we repeated the analysis using only 336 genes conserved in more than 10 species of **both** Fungi and Metazoa. Each dot represents an iteration out of 100. A root between Fungi and all other eukaryotes is the second most parsimonious. The smaller subset of genes in comparison with the original SEL+ dataset (2,786 genes) is likely more susceptible to produce artifacts by highly underrepresented taxa from clades such as Apusozoa, Glaucophytes and Malawimonas (the first, third and fifth most parsimonious hypotheses).
