## Supplementary material for "Phylogenomic Analyses Of 2,786 Genes In 158 Lineages Support a Root of The Eukaryotic Tree of Life Between Opisthokonts (Animals, Fungi and Their Microbial Relatives) and All Other Lineages": Figure S4

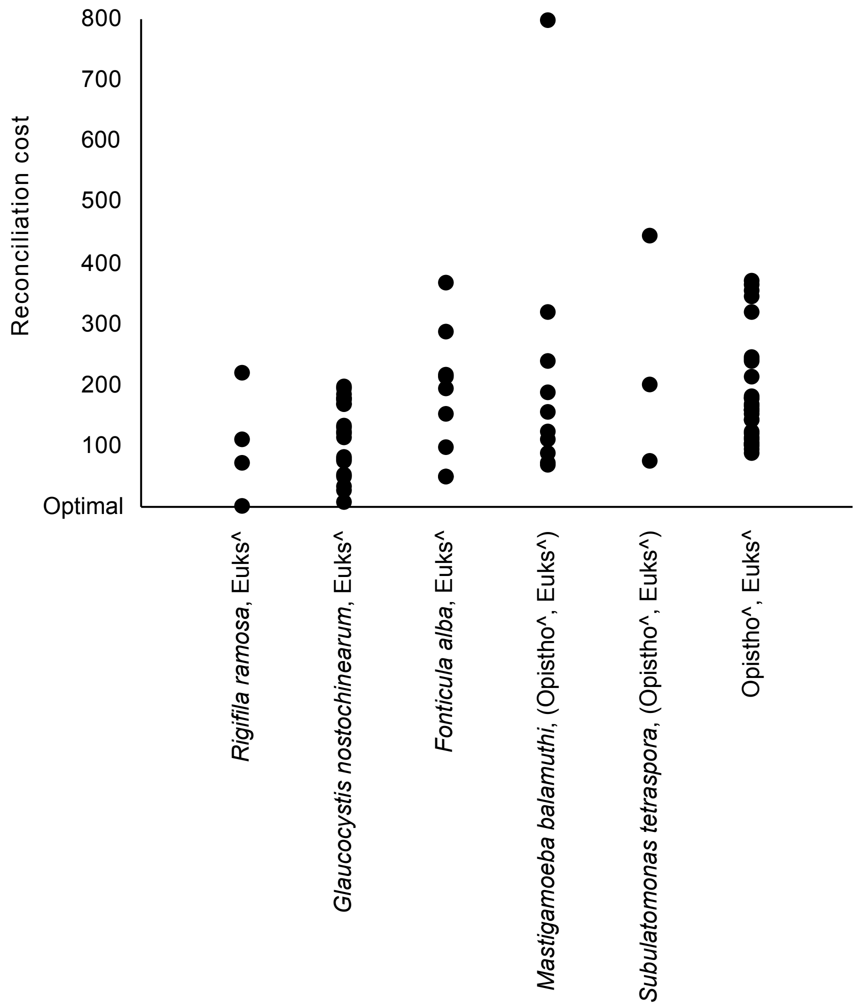


**Figure S4.** To assess whether the reduced gene numbers within fungal genomes drives our estimate of the root, we also repeated the analysis using 246 genes conserved in more than 10 species of Metazoa and absent in Fungi. Despite the lower power to detect phylogenetic signal and the reduced set of Opisthokonts (no Fungi), a root between Opisthokonta and others is one of the most parsimonious (reconciliation cost and number of iterations). As in Figure S1, the smaller dataset allows highly underrepresented clades (Apusozoa, Archamoebae, Microsporidia and glaucophytes) to impact the results, and there is less difference among reconciliation costs.
